# Privatization of a breeding resource by the burying beetle *Nicrophorus vespilloides* is associated with shifts in bacterial communities

**DOI:** 10.1101/065326

**Authors:** Ana Duarte, Martin Welch, Josef Wagner, Rebecca M. Kilner

## Abstract

It is still poorly understood how animal behaviour shapes bacterial communities and their evolution. We use burying beetles, *Nicrophorus vespilloides*, to investigate how animal behaviour impacts the assembly of bacterial communities. Burying beetles use small vertebrate carcasses as breeding resources, which they roll into a ball, smear with antimicrobial exudates and bury. Using high-throughput sequencing we characterize bacterial communities on fresh mouse carcasses, aged carcasses prepared by beetles, and aged carcasses that were manually buried. The long-standing hypothesis that burying beetles ‘clean’ the carcass from bacteria is refuted, as we found higher loads of bacterial DNA in beetle-prepared carcasses. Beetle-prepared carcasses were similar to fresh carcasses in terms of species richness and diversity. Beetle-prepared carcasses distinguish themselves from manually buried carcasses by the reduction of groups such as Proteobacteria and increase of groups such as Flavobacteriales and Clostridiales. Network analysis suggests that, despite differences in membership, network topology is similar between fresh and beetle-prepared carcasses. We then examined the bacterial communities in guts and exudates of breeding and non-breeding beetles. Breeding was associated with higher diversity and species richness. Breeding beetles exhibited several bacterial groups in common with their breeding resource, but that association is likely to disappear after breeding.

## Introduction

Advances in sequencing technology have revealed the full extent of bacterial diversity living within and alongside animals. Bacteria have been shown to play important roles in animal development, ecology and evolution (McFall-Ngai *et al.*, 2013; Ezenwa *et al.*, 2012). Likewise, animals acting as ecosystem engineers (*sensu* Jones *et al.*, 1994) likely change the biotic and abiotic conditions where bacterial communities develop. For example, disturbances caused by soil foraging animals change the composition of soil microbial communities in an arid ecosystem, leading to functional shifts that may affect nutrient cycles (Eldridge *et al.*, 2015); bioluminescent squid which harbour the bacterium *Vibrio fischeri* expel the excess cells of the symbiont daily into the surrounding water, driving abundance patterns of *V. fischeri* in marine communities (Lee and Ruby, 1994); in laboratory populations of *Drosophila* flies, the presence and density of flies changes the microbial communities in the flies’ food (Wong *et al.*, 2015). In these examples, the animals are not selected to change their bacterial environment; rather the changes in bacterial communities are a by-product of other animal adaptations.

In many other cases, however, animals are under direct selection to control or manipulate their external microbial communities; for example, fungus-growing ants and termites have special glands that secrete antibacterial and antifungal substances to protect their symbiotic fungus from competition from other microbes (Do Nascimento *et al.*, 1996; Rosengaus *et al.*, 2004; Cremer, Sophie A O Armitage, *et al.*, 2007); wood ants bring tree resin back to their nests to protect the colony against pathogens (Chapuisat *et al.*, 2007); túngara frogs protect their nests with antimicrobial proteins (Fleming *et al.*, 2009).

Another example of animals being selected to control their bacterial environment are those depending on rapidly decaying resources such as cadavers and fallen fruit. There is selective pressure on these animals to evolve behavioural and chemical defences to minimize competition with bacteria (Janzen, 1977). Supporting this view, a range of antimicrobial peptides (AMPs) and lysozymes have been found in secretions of species such as blowflies, hide beetles, mealworm beetles, and burying beetles (Barnes *et al.*, 2010; Kerridge *et al.*, 2005; Cotter *et al.*, 2010; Degenkolb *et al.*, 2011).

Burying beetles, *Nicrophorus vespilloides*, rear their offspring in small vertebrate carcasses which they prepare in a series of steps. First, they shave the fur or feathers and make a small incision in the abdomen. Then the intestines of the cadaver are pulled out and consumed (Pukowski, 1933). This is a highly repeatable behaviour across individuals; we have video-recorded it in 20 separate breeding events under laboratory conditions (C.M. Swannack, personal communication; see video file Supplementary Material). Consuming the gut is potentially crucial to avoid putrefaction caused by enteric microbes, which would lead to rupture of the body cavity and leaking of body fluids and metabolic by-products, which promotes succession on the carcass (Metcalf *et al.*, 2013). The beetles then roll the carcass into a ball, coating it with oral and anal exudates and burying it in the soil (Pukowski, 1933; Scott, 1998). By preparing the carcass in this way, the beetles are thought to be privatizing their breeding resource, because they exclude others from using it (Strassmann and Queller, 2014). Burial hides the carcass from scavenging insects and vertebrates, while the oral and anal exudates are thought to eliminate microbes (Rozen *et al*. 2008). Evidence from previous studies suggests that the function of the burying beetle’s anal exudates is to reduce bacterial load, promoting larval survival (Arce *et al.*, 2012) while decreasing both competition with bacteria and the risk of disease (Rozen *et al.*, 2008; Cotter *et al.*, 2010). When breeding on a carcass, expression of insect lysozyme and AMPs is upregulated both in the adult beetles’ gut and in the anal exudates (Cotter *et al.*, 2010; Arce *et al.*, 2012; Palmer *et al.*, 2016; Jacobs *et al.*, 2016). The effects of these antimicrobial substances against single bacterial species have been well demonstrated in a laboratory setting (e.g. Hoback *et al.*, 2004; Cotter *et al.*, 2010; Arce *et al.*, 2012). However, the effect of the beetles’ antimicrobial defences on the whole bacterial community on the carcass is unknown, and the structure of the bacterial community on the prepared carcass has never before been described. Therefore it is unclear whether burying beetles are excluding bacteria from their carrion breeding resource, or engineering a different bacterial ecosystem.

Here we study the effect of carcass preparation on the bacterial communities living in these cadavers. We ask whether bacterial load is indeed reduced when burying beetles prepare a carcass, and how the bacterial community changes in diversity and membership when beetles are present. We analyse microbial networks to investigate whether associations between bacterial groups change when beetles are present. We also characterize the bacterial communities present in the guts and anal exudates of breeding and non-breeding beetles, to describe how breeding changes the beetle’s bacterial communities and determine whether associations formed on carcasses between beetles and specific bacterial groups are limited to the duration of the breeding event, or last beyond it.

## Methods

### Bacterial communities on mouse carcasses

In September 2012, we collected *Nicrophorus vespilloides* individuals from traps at different locations in Byron’s Pool nature reserve in Grantchester, Cambridgeshire, UK (ordnance survey grid reference TL436546). Beetles were kept under standard laboratory conditions in individual boxes (12 × 8 × 2 cm) filled with moist garden compost and fed approximately 1 mg minced beef twice per week. It is not possible to accurately assess prior breeding experience or age in field-collected individuals. To ensure all experimental individuals were sexually mature and had experienced similar conditions prior to the experiment, we allowed them to breed once under standard laboratory conditions: pairs of males and females were placed in plastic breeding boxes (17 × 12 × 6 cm) half-filled with moist garden compost and provided with a thawed mouse carcass (12-16 g). After reproduction, adults were kept for use in the experiment described next.

The following week, we collected soil at six separate locations in the field near our beetle traps in Byron’s Pool. Some of the soil was placed in sterile sample bags and frozen at −80°C within 4h of collection for assessment of bacterial communities. The remaining soil was brought back to the laboratory.

We filled 19 breeding boxes to half of their height using soil collected in the field. We placed a thawed mouse carcass (Livefood UK Ltd, previously kept at −20°C) in each box and closed the lid (day 0 of the experiment). These carcasses were then subjected to one of the following three treatments. On day 1, we sampled the bacterial community on 6 of the carcasses (hereafter called the ‘fresh’ treatment). On the same day, we introduced a pair of field-caught burying beetles to 7 other carcasses (‘beetle’ treatment). Burying beetles prefer breeding in fresh carcasses over decaying ones, potentially because decayed carcasses yield them lower reproductive success (Rozen *et al.*, 2008). Hence our choice of introducing beetles after 1 day of decomposition is reflective of the study species’ natural behaviour. The remaining 6 carcasses were manually buried (approximately 2 cm deep) in the soil, to mimic the burial performed by beetles (‘buried’ treatment). On day 3, we removed beetle pairs from the beetle treatment because carcass preparation was complete. On day 4 of the experiment, we sampled the bacterial community on the beetle and buried carcasses.

To sample the bacterial communities on carcasses, we first removed as much soil debris as possible with sterile tweezers. To maximize the amount of bacterial DNA collected from every region of the carcass, we rolled the carcass in 40 ml of sterile phosphate-buffered saline (PBS) on Petri dishes, using a sterile swab to release as much material as possible from the carcass into the PBS solution. We pipetted the solution to a 50ml tube and pelleted the bacterial cells and debris at 3930 × g for 10 minutes. We discarded the supernatant and stored the pellet at −80°C until DNA extraction.

By analysing bacterial samples from carcasses in the fresh treatment, we were able to characterize the bacterial communities present on the surface of the carcass before introduction of beetles. By comparing bacterial communities on the beetle and buried carcasses, we could account for changes in microbial community that were due to carcass age and burial.

### Bacterial communities in the burying beetle’s gut and exudates

Next we analysed the bacterial communities associated with female burying beetles. We focused on females because they remain longer with the brood than males (Scott, 1998; De Gasperin *et al.*, 2015), invest more than the male in antibacterial defences (Cotter and Kilner, 2010) and provide most of the direct care (Smiseth and Moore, 2004). We collected individuals from Byron’s Pool in June 2013. Female beetles were kept under standard laboratory conditions and allowed to breed once with laboratory stock beetles, one week before the experiment took place. Hence all females were sexually mature and had bred previously, just as in the experiment analysing the carcass bacterial communities.

After breeding, females were retained. Five of those females were placed with a virgin male from the laboratory stock population in breeding boxes half-filled with moist compost and provided with a thawed mouse carcass. Three days later, at the time of larval hatching, we collected anal exudates from the breeding females, using a standard procedure we have described before (Cotter & Kilner 2010). Beetles readily produce a brown exudate when tapped gently on their abdomen. Exudates were collected with a capillary tube and diluted in 200 µl sterile PBS. On the same day, we collected exudates of four non-breeding females that had been kept in individual boxes. Breeding and non-breeding females were then anesthetized with CO_2_ and their gut was resected. Beetle guts were placed in centrifuge tubes with sterile PBS. Gut and exudate samples were stored at −80°C until DNA extraction.

### Molecular analysis

DNA was isolated using the FastDNA® Spin Kit for Soil (MP Bio Laboratories, Inc. Carlsbad, CA, USA) and stored at −20°C until use. In order to obtain an approximate measure of bacterial abundance in different carcass treatments, we performed quantitative real-time polymerase chain reaction (qRT-PCR) on a fragment of the 16S rRNA-encoding gene (detailed methods in Supplementary Material). For library construction, we first PCR-amplified the full length bacterial 16S rRNA-encoding gene (using primer pair 27F/ U1492R; Weisburg *et al.*, 1991). We confirmed the presence of the amplicon by agarose gel electrophoresis. The amplicon band was excised from the gel and the DNA was extracted with the Wizard® SV Gel and PCR Clean-Up System (Promega). DNA was quantified using a NanoDrop ND-1000 Spectrophotometer. To amplify the V3 region of the 16S rRNA-encoding gene, a second PCR was run using 10 ng of amplicon template DNA per sample, with Illumina-compatible primers and PCR conditions described in Bartram *et al*. (2011). Amplification of the V3 region was confirmed by agarose gel electrophoresis and DNA was extracted from the corresponding band (300 bp). High-throughput paired-end sequencing was carried out using an Illumina MiSeq instrument at the DNA Sequencing Facility (Department of Biochemistry, University of Cambridge). Sequence reads (accession number XXXX) were analysed using MOTHUR v.1.35.1 (www.mothur.org) software package (Schloss *et al.*, 2009), following the Standard Operating Procedure described in Kozich *et al.* (2013) and MOTHUR’s Wikipedia page (http://www.mothur.org/wiki/MiSeq_SOP, accessed August 2015). The quality filtering steps are described in detail in the Supplementary Material. In brief, we trimmed sequences to reduce sequence variation due to sequencing errors and removed sequences with many homopolymers. We aligned sequences to the SILVA release 119 reference alignment and excluded those with low search scores and low similarity to the template sequences. Sequences were further de-noised during pre-clustering by clustering sequences with a difference of 2 or fewer nucleotides. Chimeric sequences were removed using UCHIME (Edgar *et al.*, 2011). The remaining sequences were taxonomically classified by comparison against the SILVA release 119 reference database. Taxonomic assignment was made at each level, given a bootstrap value greater than 80, using the Ribosomal Database Project (RDP) Classifier (Wang *et al.*, 2007). Sequences classified as Chloroplast, Mitochondria, Archaea, Eukaryota or unknown at the kingdom level were removed. Uncorrected pairwise distances were calculated between sequence reads. Sequences at a distance threshold of 0.03 were then clustered into operational taxonomic units (OTUs), using the average neighbour algorithm (Schloss and Westcott, 2011). A consensus classification for each OTU was obtained. A data matrix was generated with every OTU and the number of reads belonging to each sample assigned to each OTU. To control for differences in the number of reads obtained per sample, we used a sub-sample of the dataset in all analyses of 10760. This was chosen because the smallest number of reads in any sample was 10760, obtained in a gut tissue sample of breeding beetles.

### Statistical analysis

#### Community richness and diversity

Differences in observed richness and diversity between carcass treatments (measured using the inverse Simpson index) were tested with ANOVA. For the analysis of community richness and diversity in beetle guts and exudates, we used type of tissue (gut or exudate) and breeding condition (breeding, non-breeding) as factors in an ANOVA. The inverse Simpson index was log-transformed for the beetle guts and exudates data, because a Levene test indicated variance heterogeneity.

#### Community membership and structure

Non-metric multidimensional scaling (NMDS) with three dimensions was applied to Bray-Curtis distance matrices, to visualize distances between samples. Differences between communities were tested with PERMANOVA in R using the adonis function (vegan package, Oksanen *et al.*, 2015). The same model structure as the ANOVAs described above was used for PERMANOVA. A posteriori comparisons between levels of factors were obtained by customizing the model's contrast matrix (R script provided in the Supplementary Material).

To identify OTUs strongly associated with each treatment and combinations of treatments, we used Indicator Species Analysis in R (Cáceres and Legendre, 2009). This is a standard community ecology approach that takes into account both relative abundance and relative frequency of occurrence in various sites (Dufrêne and Legendre, 1997). The Indicator Value is highest when all occurrences of an OTU are found in a single group of sites (i.e., treatments) and when the OTU occurs in all instances of that group (i.e., samples within a treatment).

In order to gain insight into the associations between OTUs in different carcass treatments, we used network analysis. The degree of association between OTUs within each group of samples was measured as Pearson's correlation coefficient. The OTU table was separated by groups ('fresh', 'beetle', and 'buried') and OTUs that were not present in at least 50% of the samples within each group were removed. Pearson's correlation coefficients were calculated with the rcor.test function from the ltm package in R (Rizopoulos, 2006). *P*-values were generated maintaining the false discovery rate below 5% using the Benjamini-Hochberg procedure. We considered all significant Pearson correlation coefficients to be network edges. Visualization of these networks was done using the R package igraph (Csardi and Nepusz, 2006), incorporating taxonomic data (see Supplementary Material). OTUs without significant correlations with other OTUs were not included in calculations of topological metrics. We calculated topological metrics of density and connectivity for each network in Cytoskape version 3.3.0 for Unix (www.cytoskape.org), using the network analysis tool.

## Results

### Bacterial communities on the carcass

#### Bacterial DNA concentration

There was an overall effect of carcass treatment on bacterial DNA concentration (Kruskal-Wallis test: X^2^ = 7.15, *p* = 0.03; Figure 1). Beetle-prepared carcasses had significantly higher concentrations of bacterial DNA than fresh carcasses (Dunn test: *Z* = 2.37, Bonferroni-adjusted *p* = 0.026) and manually buried carcasses (*Z* = −2.15, Bonferroni-adjusted *p* = 0.047). Bacterial DNA concentration did not differ significantly between fresh and buried carcasses (*Z* = 0.10, *p* = 1.0).

**Figure 1.**
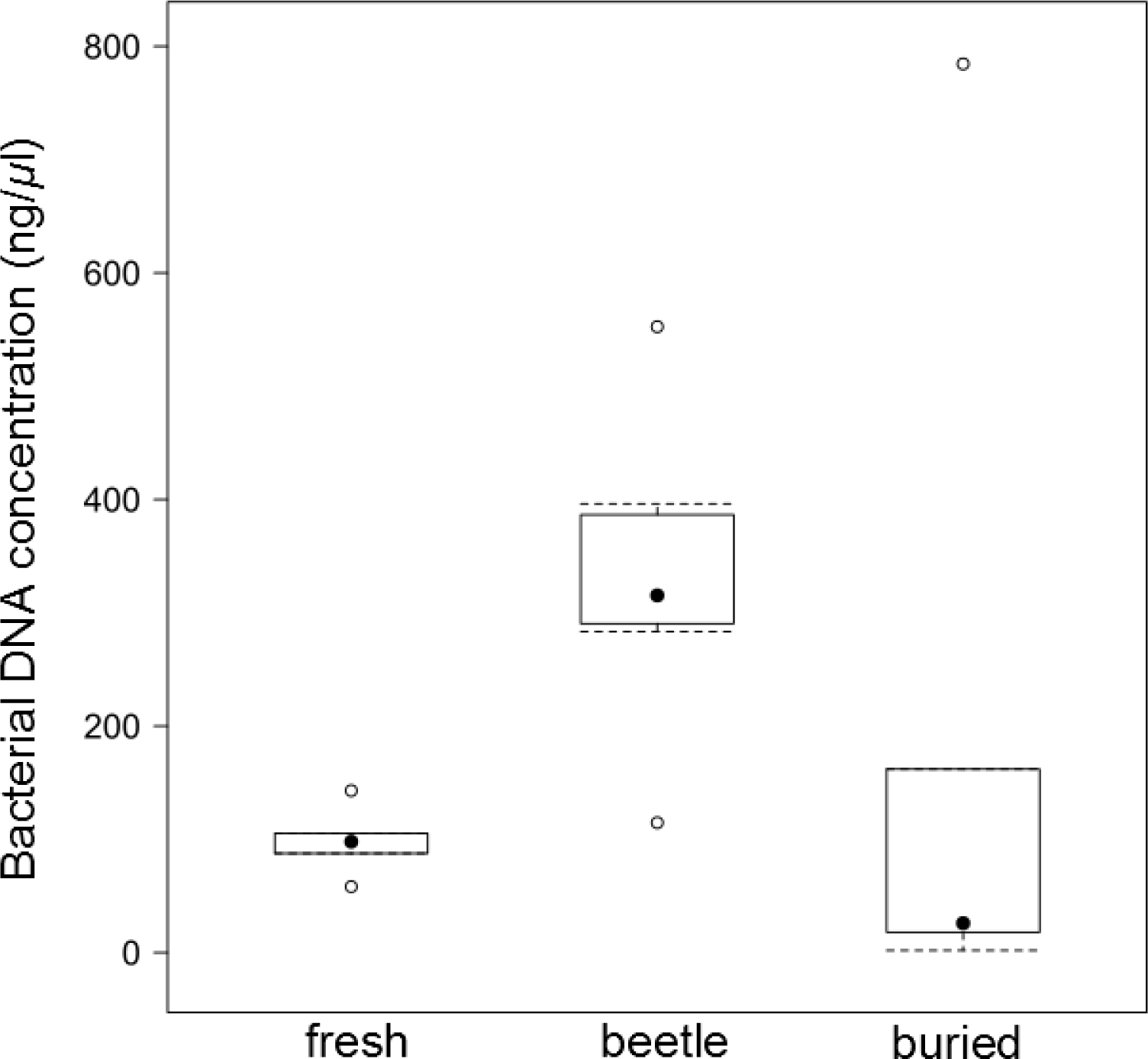
Concentration of bacterial DNA (ng/µl) quantified by quantitative real-time PCR in the three carcass treatments: fresh, beetle-prepared, and buried.

#### Richness and α-diversity of bacterial communities

Across all samples we found that most OTUs occurred at very low sequence abundances. After subsampling, the OTU table for soil samples contained 2653 OTUs, with just 9% of those responsible for 80% of total sequence abundance. In carcass samples, after subsampling, 549 OTUs remained. Just 7 of those OTUs (1.4% of the total) contributed to 80% of the total sequence abundance.

The highest OTU richness and diversity was observed in soil samples (Table 1). Observed richness was significantly higher in buried carcasses than in fresh and beetle-prepared carcasses (Table 1; effect of treatment: *F*_2_ = 10.62, *p* = 0.001). Carcasses prepared by beetles and fresh carcasses showed similar levels of observed richness (Table 1; Tukey post-hoc comparison: adjusted *p* = 0.76). Bacterial taxonomic diversity, assessed as the inverse Simpson index, also differed significantly between carcass treatments (effect of treatment: *F*_2_ = 5.93, *p* = 0.012). Buried carcasses showed the highest diversity, while beetle-prepared and fresh carcasses showed similar levels of diversity. Diversity was higher in buried than fresh (adjusted *p* = 0.012) and beetle carcasses, although in the latter case the difference was marginally non-significant (adjusted *p* = 0.087). Fresh and beetle-prepared carcasses showed similar levels of diversity (adjusted *p* = 0.451). Hence, despite the higher concentration of bacterial DNA, indicating that beetle carcasses are a prolific ground for bacteria, beetle-prepared carcasses exhibited relatively fewer species than unprepared carcasses of the same age.

**Table 1.** Means and standard errors of observed richness and diversity (inverse Simpson) index for the different sample types.

| Type of sample | Observed richness | Inverse Simpson index |
| --- | --- | --- |
| Soil | 1078.95 ± 36.07 | 73.65 ± 7.24 |
| Fresh carcass | 62.97 ± 5.32 | 1.76 ± 0.45 |
| Beetle carcass | 50.83 ± 12.25 | 2.59 ± 0.40 |
| Control carcass | 126.48 ± 17.73 | 4.13 ± 0.60 |
| Beetle guts – breeding | 79.22 ± 6.74 | 3.39 ± 0.87 |
| Beetle guts – non-breeding | 39.24 ± 4.69 | 1.16 ± 0.03 |
| Beetle exudates – breeding | 73.66 ± 4.64 | 3.43 ± 0.32 |
| Beetle exudates – non-breeding | 65.39 ± 9.73 | 1.80 ± 0.36 |

#### Community composition

There were significant differences in bacterial community composition across all treatments. Soil communities differed significantly from carcass communities (Pseudo *F* = 10.554, *P*(Perm) = 0.001; Figure 2). Communities on fresh carcasses were significantly different from communities on beetle-prepared carcasses (Pseudo *F* = 9.167, Bonferroni-corrected *P* = 0.002). There were also significant differences between beetle-prepared and control carcasses (Pseudo *F* = 6.071, Bonferroni-corrected *P* = 0.002).

**Figure 2.**
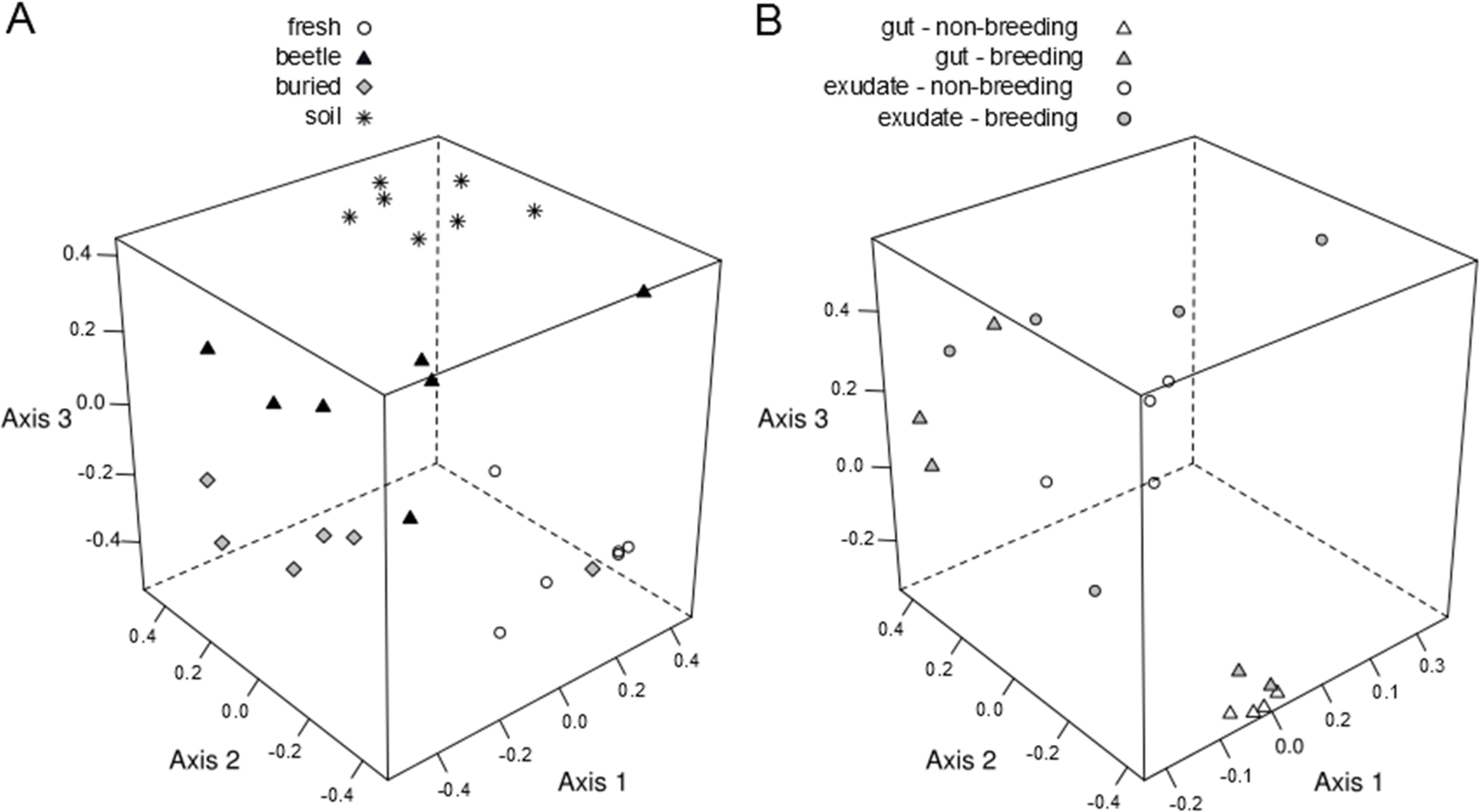
Non-metric multi-dimensional scaling plot of the three dimensions of an ordination of **A**) 2653 bacterial OTUs present in soil and carcass samples, and **B**) 284 bacterial OTUs in beetles’ gut and exudates.

The most common bacterial phyla in soil communities were Actinobacteria and Proteobacteria (Figure 3a). Acidobacteria and Bacteroidetes were also common, yet contributed lower proportions of reads. There were also 472 OTUs that could not be classified to phylum level.

**Figure 3.**
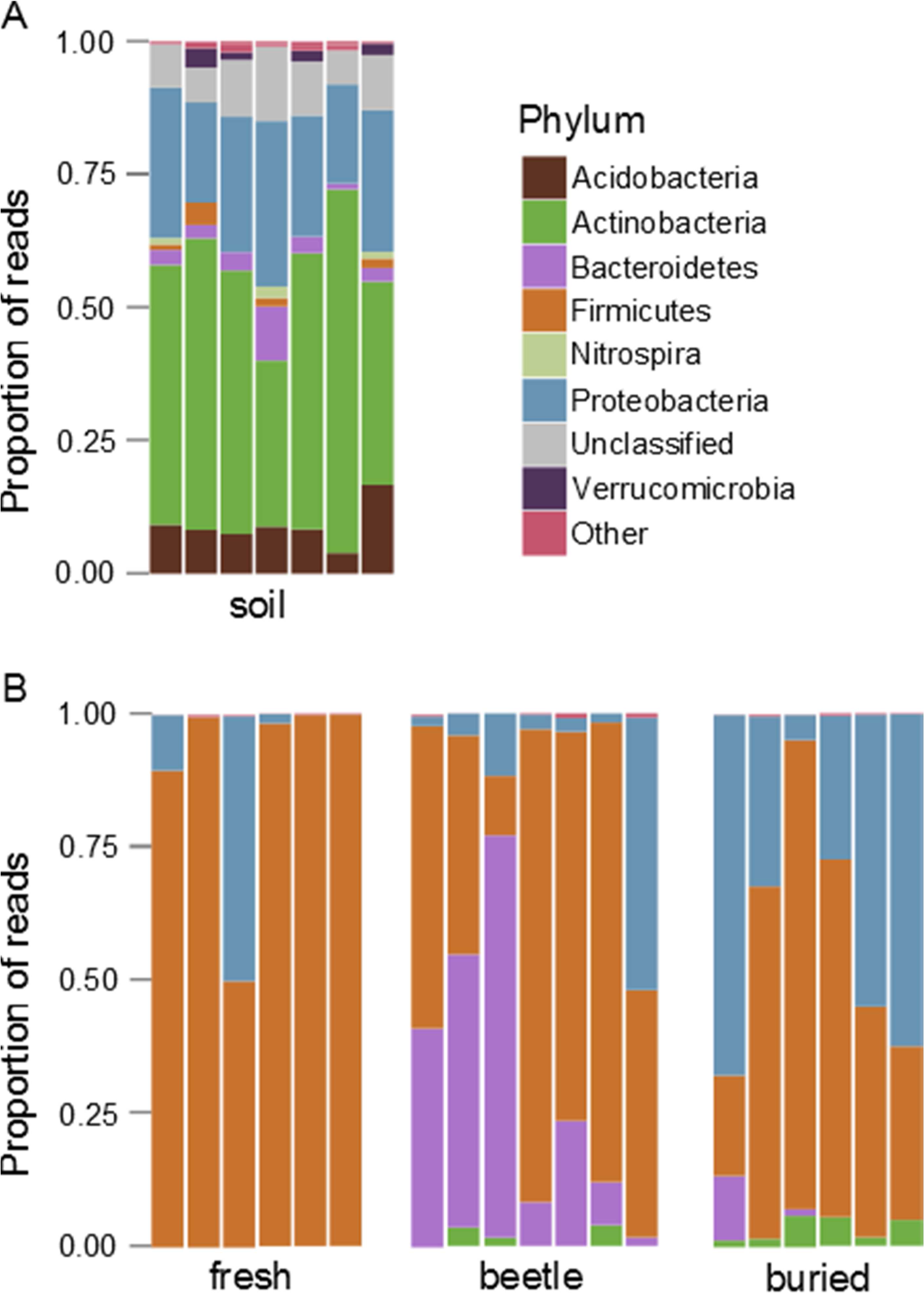
Relative abundance of major bacterial taxa within **A**) soil samples and **B**) carcass samples. Each bar represents a different sample. Carcass samples are ordered to match the soil samples corresponding to the original soil on which carcasses were laid.

In fresh carcasses, the majority of reads belonged to Firmicutes OTUs (Figure 3b). In one sample, OTUs belonging to Proteobacteria composed approximately 50% of the observed reads, but this phylum was observed in low proportions in all other samples. Beetle-prepared samples were mostly comprised of Firmicutes, Bacteroidetes and, in one sample, Proteobacteria. Manually buried carcasses were mostly dominated by Firmicutes and Proteobacteria. The finding that Firmicutes and Proteobacteria are the most common phyla in 4-day old carcasses is in agreement with previous characterizations of microbial communities in mouse cadavers (Metcalf *et al.*, 2013).

#### Indicator analysis

We have seen that bacterial communities differ between carcass types. To understand what groups are driving these differences, we used Indicator Species Analysis to identify OTUs significantly associated with different types of samples. Just three Bacillales OTUs were indicator species of fresh carcasses. For beetle-prepared carcasses, one Microbacteriaceae, one Planococcaceae, two Clostridiales (one unclassified and one *Tissierella*), two Flavobacteriaceae (one unclassified and one *Myroides*), two Enterococcaceae and one Moraxellaceae (*Acinetobacter*) OTU were indicator species. For buried carcasses, one Micromonosporaceae, two Planococcaceae, one Enterobacteriaceae (*Escherichia-Shigella*), two Pseudomonadaceae (*Pseudomonas*), and one unclassified Alphaproteobacteria were indicator species (Table 2).

**Table 2.**
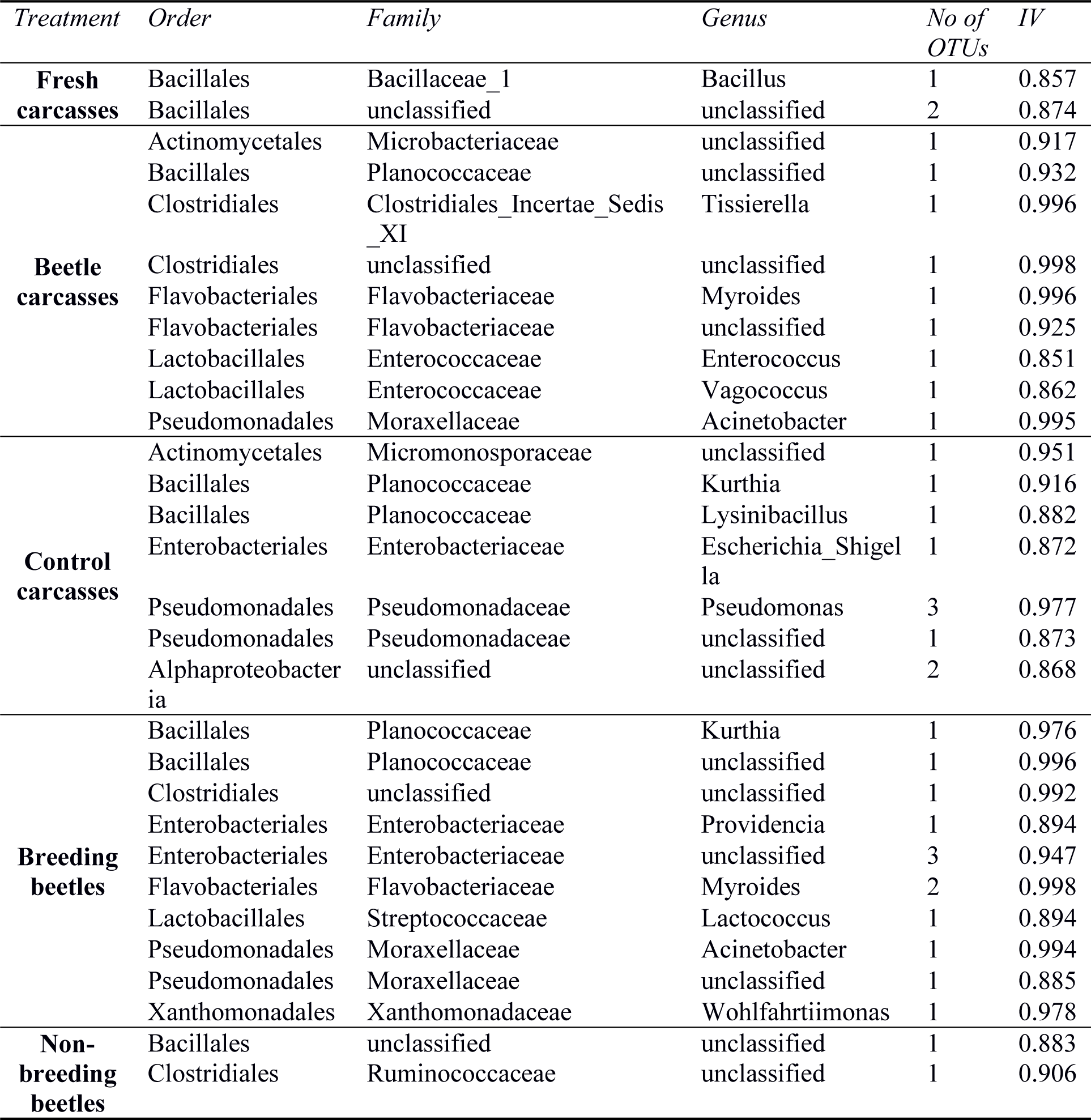
Bacterial taxa associated with different treatments using Indicator Species Analysis. Gut and exudates samples were grouped by breeding status to identify indicator species of breeding versus non-breeding beetles. Only significant (p < 0.05) taxa with Indicator Value (IV) > 0.85 are shown.

#### Network analysis

Network analysis has been used to help form hypotheses on the relationships between members of microbial communities (Chaffron *et al.*, 2010; Barberan *et al.*, 2012). Here we examined correlations in OTU abundances within carcass treatments to gain insight on how bacterial groups co-occur in the different environments.

Network visualizations are shown in the Supplementary Material. The number of vertices was similar among fresh, beetle-prepared and buried carcasses (Table 3), despite the higher OTU richness in buried carcasses. Across all treatments, significant correlations between OTUs were positive, rather than negative. Network connectivity for all treatments was zero, i.e. there was no path connecting every vertex in the networks. This resulted in separate clusters of connected OTUs (see Supplementary Material). The highest number of edges (i.e. significant correlations), mean number of neighbours, clustering coefficient and density was found in fresh carcasses (Table 2). Network metrics for beetle-prepared carcasses showed slightly lower values than network metrics of fresh carcasses, with exception of network centralization. In comparison, manually buried carcasses showed the fewest significant correlations between OTUs and the lowest values for all network metrics, indicating that the relationships between OTUs observed in fresh carcasses breakdown over the course of decomposition.

**Table 3.**
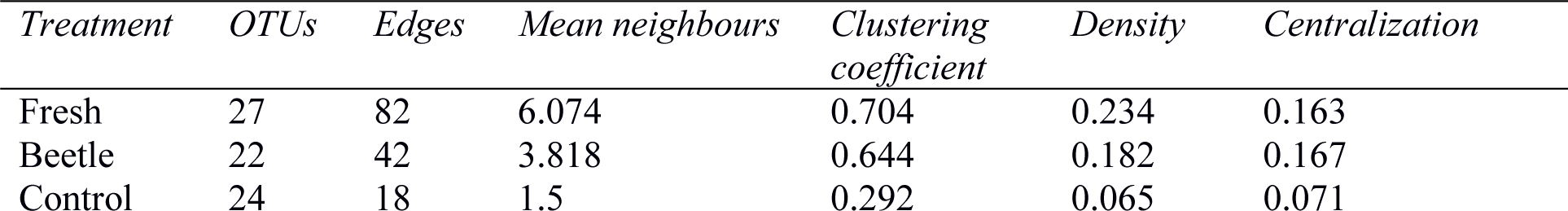
Metrics for networks of bacterial communities in carcass samples.

| *Treatment* | *OTUs* | *Edges* | *Mean neighbours* | *Clustering*<br>*coefficient* | *Density* | *Centralization* |
| --- | --- | --- | --- | --- | --- | --- |
| Fresh | 27 | 82 | 6.074 | 0.704 | 0.234 | 0.163 |
| Beetle | 22 | 42 | 3.818 | 0.644 | 0.182 | 0.167 |
| Control | 24 | 18 | 1.5 | 0.292 | 0.065 | 0.071 |

### Bacterial communities in N. vespilloides guts and exudates

#### Richness and α-diversity of bacterial communities

Again, most OTUs occurred at very low sequence abundances: 284 OTUs remained after subsampling, of which just 4 (1.4% of the total) contributed to more than 80% of total sequence abundance. OTU richness was overall higher in breeding beetles than in non-breeding beetles (Table 1; *F* = 13.22, *p* = 0.003). The lowest observed richness was found in guts of non-breeding beetles. We found a significant interaction between breeding status and type of tissue (gut or exudate), largely driven by a difference in richness between guts and exudates of non-breeding beetles, although statistically, this difference was marginally non-significant (*p* = 0.080). Breeding beetles showed higher diversity than non-breeding beetles (*F* = 12.25, *p* = 0.003), independent of the type of tissue (guts or exudates).

#### Community composition

Breeding status had a significant effect on the composition of bacterial communities (Pseudo-F = 5.189, *P*(Perm) = 0.016). This suggests that the changes in bacterial communities that arise when beetles are breeding are temporary and environmentally induced. We again found an interaction between breeding status and type of tissue (*P*(Perm) = 0.010). To further investigate this, we customized model contrasts for post-hoc testing (Supplementary Material). The model with customized contrasts revealed that differences between communities in guts and exudates of breeding beetles were marginally non-significant (*P*(Perm) = 0.086), whereas in non-breeding beetles, guts and exudates showed significant differences (Pseudo-*F =*10.771, *P*(Perm) = 0.001; Figure 4). A possible reason for this is that guts of non-breeding beetles were very homogenous, dominated by Firmicutes (mostly *Bacillus* spp.) in all samples (Figure 4a). Exudates of non-breeding beetles show more variability between samples, perhaps because exudates come into contact with bacteria in the beetle’s external cuticle. In breeding beetles, both gut and exudate samples show variability, which makes it statistically difficult to differentiate between communities. As was found in beetle-prepared carcasses, exudates from breeding beetles showed a high proportion of Bacteroidetes (Figure 4b). Bacteroidetes were also present in three gut samples in breeding beetles. Proteobacteria and Actinobacteria also comprised a small proportion of reads in breeding beetles.

**Figure 4.**
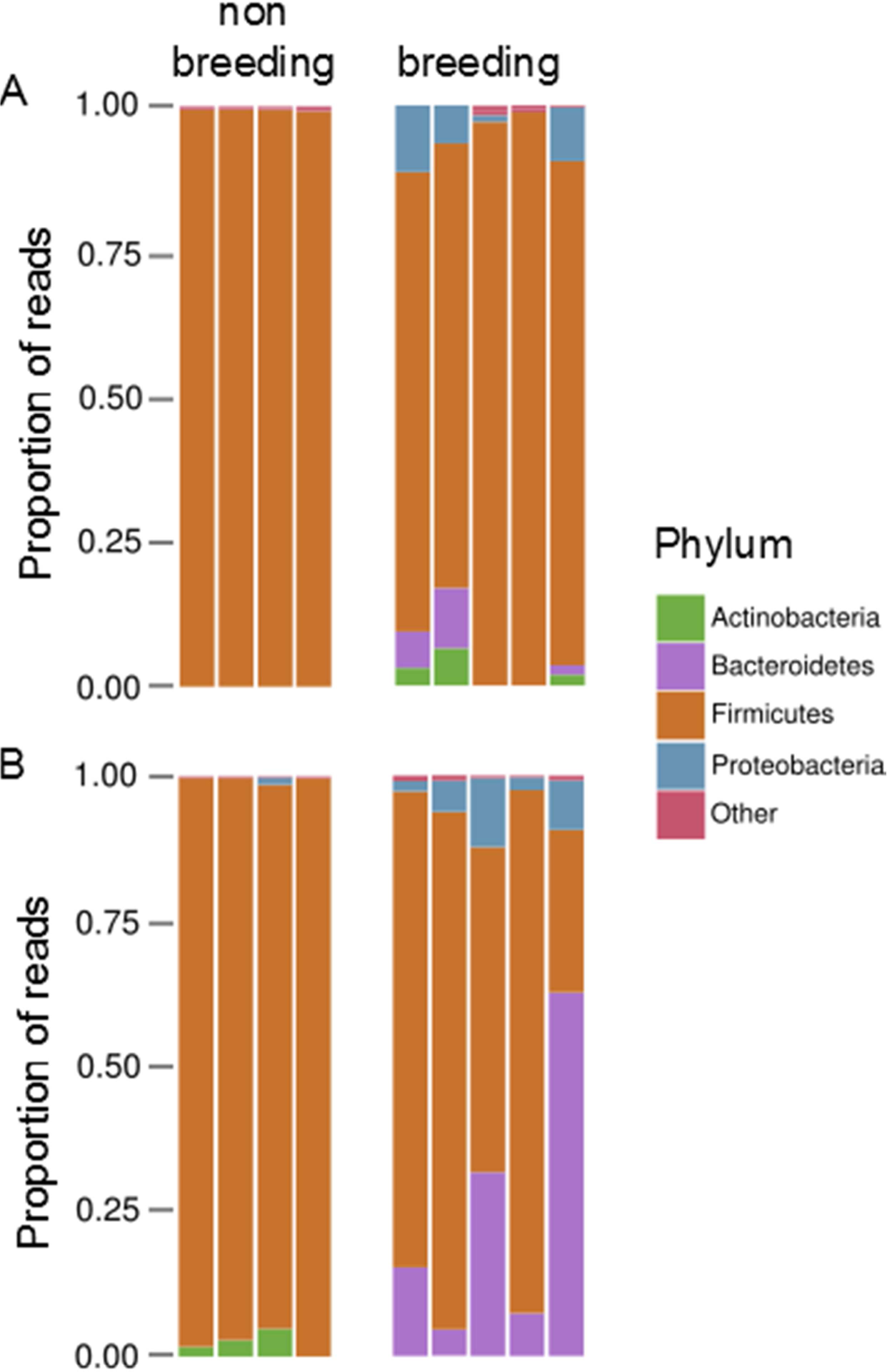
Relative abundance of major bacterial taxa within **A**) gut of non-breeding and breeding beetles, and **B**) exudates of non-breeding and breeding beetles.

#### Indicator analysis

We searched for indicator species that distinguish communities in breeding beetles from communities in non-breeding beetles, combining tissues within those categories. One unclassified Bacillales and one Ruminococcaceae (Clostridiales) OTU were the only indicators of communities of non-breeding individuals. Breeding beetles shared four indicator OTUs with beetle-prepared carcasses: one Flavobacteriaceae (*Myroides*), one unclassified Planococcaceae, one Moraxellaceae (*Acinetobacter*) and one unclassified Clostridiales OTU. Breeding beetles shared a single Planococcaceae OTU (*Kurthia*) with buried carcasses. Furthermore, four Enterobacteriaceae, one Streptococcaceae (*Lactoccocus*), one unclassified Moraxellaceae and one Xanthomonadaceae (*Wohlfartiimonas*) OTU were indicators of breeding beetles, in comparison to non-breeding beetles.

## Discussion

The impact of animal behaviour on microbial communities in the environment is still poorly understood. Here we show that burying beetles, which use small vertebrate carcasses as breeding resources, change the bacterial communities living on those carcasses. We found that burying beetles increase bacterial load when they prepare a carcass, and that there are substantial differences in richness, diversity and composition of bacterial communities developing on beetle-prepared and unprepared carcasses. Bacterial communities in burying beetle guts and exudates showed similarities with the carcasses upon which they breed, but these resemblances were confined to the breeding event, indicating transient associations between beetles and environmental bacteria.

Our first finding refutes the current assumption that burying beetles ‘clean’ and reduce the bacterial load on their carcasses (e.g. Rozen *et al.*, 2008). On the contrary, our finding reveals that carcass preparation (of relatively fresh carrion) is associated with an increase in bacterial load, rather than a reduction. Our results support the conclusions of a recent behavioural study examining resource use in larvae of two species closely related to *N. vespilloides*, which suggests that the application of antimicrobial exudates on carcasses has not evolved primarily as a response to microbial competition for carrion (Trumbo *et al.*, 2016). Our study did not set out to test concurrent hypothesis for the evolution of antimicrobial exudates in burying beetles, yet the absence of several known pathogens in beetle-prepared carcasses lends support to the hypothesis that antimicrobial exudates serve as social immunity against pathogens (Cremer *et al.*, 2007; Cotter and Kilner, 2010).

Despite the increased bacterial load, we found that bacterial communities on beetle-prepared carcasses showed levels of species richness and diversity as low as those associated with fresh carcasses, and lower than on the carcasses we manually buried. One explanation could be that the buried carcasses were colonized by soil bacteria, which are highly diverse (Figure 2a). However, beetle-prepared carcasses were also buried in the soil yet exhibited less bacterial diversity. We suggest that the antimicrobial substances produced by beetles restrict the diversity of species that can grow on carcasses, reducing the competition for resources and thus favouring the growth of species that beetles do not eliminate.

Consistent with this idea, community membership differed between beetle-prepared carcasses and both fresh and manually buried carcasses (Figure 3b). In part, this was due to the selective elimination of some groups of bacteria by burying beetles. We expected to see a higher proportion of Gram-bacteria on beetle-prepared carcasses, because a key antibacterial compound present in burying beetle exudates is lysozyme (Cotter *et al.*, 2010; Jacobs *et al.*, 2016), which is mainly effective against Gram+ bacteria. We found support for this prediction when comparing the bacterial communities found on fresh carcasses with those from beetle-prepared carcasses. However, comparing beetle-prepared carcasses with carcasses that we buried revealed that burying beetles are capable of eliminating some Gram-bacteria too. This is because the majority of the Gram-OTUs on beetle-prepared carcasses were Bacteroidetes, whereas on manually buried carcasses, most Gram-OTUs were Proteobacteria. Proteobacteria include several insect pathogens, such as *Serratia*, *Pseudomonas*, *Enterobacter* and *Escherichia*-*Shigella* sp. (Bulla *et al.*, 1975), hence there is potentially strong selective pressure for beetles to reduce the abundance of these bacterial groups on their breeding resource. Currently, we can only speculate about what other mechanisms are in place to control Proteobacteria on carcasses prepared by burying beetles. It is possible that burying beetles manipulate the physical conditions, such as pH, on the carcass to prevent the growth of several common Proteobacteria (the pH of anal exudates is approximately 9, A.D., personal observation). Alternatively, burying beetles may promote colonization of the carcass by other bacteria, which then eliminate Proteobacteria.

Community composition on burying beetle-prepared carcasses was also changed by the addition of some groups of bacteria. The presence of beetles on a carcass was associated with Clostridiales, Lactobacillales and Flavobacteriales (Table 2). The first two groups are common in the mammalian gut (Madigan *et al.*, 2010). Their presence on the outside of the carcass could result from the consumption of the mouse intestine by the burying beetle and subsequent release of the mouse gut microbiota on the carcass through oral and/or anal exudates. The third group, Flavobacteriales, are a small part of the mammalian skin flora and are also present in soil and water (Madigan *et al.*, 2010). The genus *Myroides*, which was one of the most abundant Flavobacteriales in beetle-prepared carcasses (Supplementary Material), has been found in the gut of lepidopterans (Spiteller *et al.*, 2000) and flesh flies (Dharne *et al.*, 2008). The strain isolated from flesh flies shows resistance to a wide spectrum of antibiotics, as well as inhibitory effects on the growth of other bacterial species. In future work, it would be interesting to test whether these bacteria form a mutualistic relationship with the beetles by playing an active part in inter-species “bacterial warfare”. Another possibility is that they are commensals that benefit from the exclusion of other bacteria or from the changes in physical conditions on the carcasses.

We used network analysis to understand more about how bacterial groups interact with each other in the different carcass treatments. Our results suggest that bacterial communities on all types of carcasses are poorly connected and highly modular. Given that beetle-prepared carcasses have experienced a greater disturbance than control carcasses, we would expect them to show network characteristics of ‘stressed’ communities, such as reduced correlations among species (e.g. Sun *et al.*, 2013). Instead we found that the network on beetle-prepared carcasses exhibited more correlations among species than the network on manually buried carcasses and was topologically more similar to the network on fresh carcasses. Interactions among bacterial groups contribute to emergent properties of the microbiome in which they live (Ley *et al.*, 2006). Our network analyses suggest that bacterial communities on fresh and beetle-prepared carcasses might be more likely to exhibit emergent properties than those on carcasses we buried ourselves.

To understand more about the role of the burying beetle in restructuring bacterial communities, we examined the bacterial communities in gut and exudates of *N. vespilloides* females formed during and after the breeding event. Communities in breeding beetles differed from (non-virgin) non-breeding beetles (Figure 4), as might be expected given the changes in diet, social environment and physiological state when beetles breed. One key finding was that bacterial species richness and diversity was lower in the guts of non-breeding females than the guts of breeding females. This may be because some aspects of the internal immune system are down-regulated during breeding (Cotter *et al.*, 2013; Reavey *et al.*, 2014; Jacobs *et al.*, 2016), meaning that greater abundances of gut bacteria are picked up from the environment. Perhaps non-breeding individuals are more capable of keeping environmental bacteria out of their digestive tract than breeding beetles.

A second key finding was that breeding beetles have several indicator species in their guts and exudates in common with communities on beetle-prepared carcasses (Table 2). Furthermore, these bacteria are generally absent or low in abundance in non-breeding beetles. This indicates that associations between beetles and particular bacterial OTUs are transient, are promoted by their association with the breeding resource and are unlikely to last beyond the breeding event.

A previous study using high-throughput sequencing to examine microbial communities in the hindgut of Silphidae beetles, found that wild-caught beetles of the genus *Nicrophorus* shared several bacterial groups (Kaltenpoth and Steiger, 2014). We found many of these groups in our samples of breeding beetles, such as Lactobacilalles (*Vagococcus* spp.), several Clostridiales, a Xanthomonadaceae (related to *Ignatzschineria* or *Wohlfartiimonas* spp), Erysipelotrichales and Neisseriales (Supplementary Material S3). Laboratory-reared *N. vespilloides* in Kaltenpoth and Steiger lacked many of those groups, which suggests that bacterial groups common to Nicrophoridae are picked up from the environment, because they feed and breed on similar species of dead vertebrate. We also found very low bacterial diversity in our non-breeding individuals, which were wild-caught but fed on minced beef before being sacrificed. This supports the conclusion that at least some of the key bacteria common to the gut microbiota of *Nicrophorus* species are environmentally acquired on their breeding resource.

## Conclusions

This study describes the effect of animal behaviour on bacterial communities in the environment where they breed. We show that the behavioural and chemical defences of burying beetles change the diversity and richness of bacterial communities on the vertebrate carcasses used as a breeding resource. Our data are not consistent with the notion that these defences rid the carcass of bacteria. Instead, they suggest that beetle defences induce a shift in bacterial community membership, and increasing the bacterial load on the carcass in general. Our analyses further suggest that carcass preparation by burying beetles involves some hitherto undescribed mechanism for elimination of potential pathogenic Gram-bacteria. The challenges for future work are to investigate the functionality of the bacterial community associated with breeding beetles, and whether the shift in bacterial community is adaptive for burying beetles or simply a by-product of carcass preparation. This work establishes the potential of the burying beetle as a system to study the manipulation and assembly of microbiotas.

## Acknowledgements

We thank Bram Kuijper for help with R code; Peter Davenport for initial data exploration; members of the Salmond and Welch groups for their help; Alecia Carter for help with network analysis; Chris Swannack for videos of carcass preparation; Amy Backhouse and Ornela De Gasperin for help in field collections; members of the Behavioural Ecology group for helpful discussions. AD was supported by NERC grant NE/H019731/1 and ERC Consolidators grant 310785 BALDWINIAN_BEETLES, both to RMK. RMK was supported in part by ERC Consolidators grant 310785 BALDWINIAN_BEETLES.

